# Temporal changes in species composition affect a ubiquitous species’ use of habitat patches

**DOI:** 10.1101/528935

**Authors:** Ellen K. Bledsoe, S. K. Morgan Ernest

## Abstract

Across landscapes, shifts in species composition often co-occur with shifts in structural or abiotic habitat features, making it difficult to disentangle the role of competitors and environment on assessments of patch quality. Using 22 years of rodent community data from a long-term experiment, we show that a small, ubiquitous granivore (*Chaetodipus penicillatus*) shifted its use of different experimental treatments with the invasion of a novel competitor, *C. baileyi*. Changes in population metrics such as residency, probability of movement between patches, and the arrival of new individuals in patches resulted in changes in which treatment supported the highest abundances of *C. penicillatus*. Our results suggest that the invasion of a new species worsened the quality of the originally preferred treatment, probably through its impacts on resource availability. Paradoxically, the invader also increased *C. penicillatus*’ use of the less preferred treatment, potentially through shifts in the competitive network on those plots.

## Introduction

Species often exist in landscapes consisting of a patchwork of habitats, some of which are conducive to a species’ survival and reproduction and others which are less suitable. Building upon intraspecific habitat selection (Fretwell & Lucas 1970) and optimal patch use theory (Charnov 1976, Brown 1988), numerous studies have shown that interspecific habitat selection can act as a potential mechanism of species coexistence (e.g., Grant 1971, Schoener 1974, Morris 1989, Morris 2003). Landscape heterogeneity creates niche opportunities (Comins & Noble 1985) while variability in patch connectivity supports scenarios where populations can be rescued from extinction by dispersal (Brown & Kodric-Brown 1977) or where species can find isolated refuges when they would otherwise be driven extinct (Sedell et al. 1990). However, changes in patch conditions can also take place through time (Ernest et al. 2008), resulting in patches that vary in their suitability for a species as conditions change. Despite the fact that population dynamics and regional processes (i.e., dispersal, colonization, extinction) are inherently both spatial and temporal, temporal variation in patches is rarely incorporated into studies on patch preference.

While many studies on habitat selection focus on differences between patches in structural habitat (e.g., vegetation structure) or abiotic conditions (e.g., temperature, soil conditions), species density and composition can also affect patch preference (Grant 1971, Morris 1989, Danielson & Gaines 1987, Abramsky et al. 1992). Understanding a species’ response to species composition is complicated because shifts in species composition often co-occur across landscapes with shifts in structural or abiotic habitat features (Whittaker 1967, Tews et al. 2004). Species may be less abundant or absent from certain patches, but whether that absence is due to the patch having an incompatible environment, the isolation of the patch from other patches, the presence of superior competitors, or a combination of any of the above reasons is often unclear. Thus, in determining how a species uses the landscape through time, it is often difficult to disentangle the roles of structural or abiotic habitat qualities from species interactions (Grant 1971, Kraft et al. 2015).

In this study, we use time-series data from a desert rodent community in southeastern Arizona, USA, to show how both spatial and temporal variation in the species composition in patches affect species and their use of habitat patches. Our system has both control plots (all rodents have access) and manipulated plots in which kangaroo rats, the behaviorally dominant genus in the system, are selectively excluded. Recent studies have shown only minimal impacts of the treatment on the plant community (Supp et al. 2012); thus, this system creates a landscape with patches of differing quality due primarily to differences in rodent species composition.

In the mid-1990s, a species of large pocket mouse (*Chaetodipus baileyi*) colonized and exhibited a preference for kangaroo rat exclosures. Here, we ask how the patch preference of a small, subdominant pocket mouse, *C. penicillatus*, across this landscape of patches with and without kangaroo rats was impacted by the arrival of the larger pocket mouse, *C. baileyi*. Because congeners are expected to compete more strongly due to their shared evolutionary history, we hypothesized that 1) *C. penicillatus* use of treatment would change with the establishment of *C. baileyi*, 2) the magnitude of change would be correlated with *C. baileyi* abundance, and 3) *C. penicillatus* residency, probability of moving between treatments, and recruitment of new individuals would show corresponding changes with the establishment of *C. baileyi*.

## Methods

### Study System and Data

We used a 22-year time series (1988 – 2010) of capture-mark-recapture rodent data collected from the Portal Project to assess how *C. penicillatus* responded to the arrival of a novel competitor, *C. baileyi*, through time. While *C. penicillatus* has been caught at the site throughout the time series, *C. baileyi* was first caught at the site in 1995. Over the next decade, *C. baileyi* became a dominant species in the system; in the late 2000s, however, *C. baileyi’s* population crashed and did not rebound (Appendix S1: Fig. S4). While *C. baileyi* individuals continue to be caught occasionally, the species is no longer dominant (see Appendix S1 for more details).

The Portal Project is a long-term experimental system (Brown 1998) located in the Chihuahuan desert near Portal, Arizona, USA, on colonized land of the Chiricahua Apache now under the jurisdiction of the U.S. Bureau of Land Management. The site consists of 24 50×50 m fenced plots with three treatments. In control plots (*n* = 10), holes cut in the fence are large enough to allow all rodent species access while full rodent removal plots (*n* = 6) have no gates. Kangaroo rat exclosures (*n* = 8) have small holes in the fences which allow passage of all rodents except for kangaroo rats (*Dipodomys* genus), which are behaviorally dominant in the system (Brown & Heske 1990).

In this study, we primarily use data from 18 plots—controls and kangaroo rat exclosures—though we include data from full removals in calculations of residency and probability of movement between treatments to increase estimate accuracy. Each plot consists of 49 evenly-spaced permanent trapping stations in a 7×7 grid. Rodent abundance and composition data are collected monthly using Sherman live traps (Ernest et al. 2018). We identify trapped individuals to species, measure and record size and sex characteristics, and give each rodent an individualizing marker (previously toe and ear tags, now exclusively passive integrated transponder (PIT) tags).

We only use data through 2010 because this is when *C. baileyi* was no longer dominant in the system. Including additional data through 2014 (treatments for many plots were changed in early 2015) produced qualitatively similar results (Appendix S1: Fig. S1-S3).

### Patch use of C. penicillatus in response to C. baileyi abundance

We assessed changes in relative abundance of *C. penicillatus* between control plots and kangaroo rat exclosures by fitting a linear model along the 1: 1 line of mean *C. penicillatus* per plot by year in kangaroo rat exclosures against control plots. We then fit a linear generalized least squares model (*nlme*, Pinheiro et al. 2018) of mean *C. baileyi* per plot by year against the resulting residuals from the previous model, accounting for temporal autocorrelation (see supplements), to evaluate how *C. penicillatus’* habitat use shifted with increases in *C. baileyi* abundance.

### Population-level metrics of C. penicillatus

For *C. penicillatus* in each treatment, we calculated apparent survival (*S*), transition probability (Ψ), and the average number of new individuals per treatment. Both survival and transition probability were estimated through a multistate capture-mark-recapture model using the *RMark* package, an R interface for the MARK software (White & Burnham 1999, Laake 2013). Different strata represented treatment types, and each time step was a trapping period. We designated each time period as being either before or after the establishment of *C. baileyi*; the first trapping period in which *C. baileyi* was caught in all eight kangaroo rat exclosures (July 1997) was used as the differentiating time point. We assumed that recapture probabilities (*p*) were equal between the treatments. Our data do not allow for differentiation between permanent emigration and death, so these two processes are not differentiated in our survival estimates; we believe that any differences in apparent survival are driven primarily by emigration and, therefore, will hereafter refer to this metric as residency. We used Program CONTRAST, a program designed specifically for comparing survival estimates (Hines & Sauer 1989), to run chi-squared tests to determine the significance of differences in residency and transition probabilities between *C. baileyi* establishment and treatments.

We calculated the number of new *C. penicillatus* individuals, defined as individuals caught and given identification tags for the first time, in each treatment. Using mean new *C. penicillatus* individuals per plot by year, we ran a linear mixed effects model *(nlme*, Pinheiro et al. 2018) to assess the interaction between treatments and *C. baileyi* establishment.

### System-level aspects of patch preference

Changes in species composition can have substantial effects on the energy use in a system (Ernest & Brown 2001). To determine how this aspect of ecosystem functioning might have contributed to *C. penicillatus* use of patches through time, we calculated the ratio of total energy use per year between the kangaroo rat exclosures and controls (*portalr*, Yenni et al. 2018).

### Analyses

All analyses were performed using R 3.5.0 (R Core Team 2018). Data and code are available on GitHub (https://github.com/bleds22e/PP_shifts).

## Results

### Patch use of C. penicillatus in response to C. baileyi abundance

After its arrival in 1995, *C. baileyi* increased in abundance until the late 2000s (Fig. 1a) and was found far more frequently on the kangaroo rat exclosures than the control plots (Appendix S1: Fig. S4). *C. penicillatus*’ preferences for the two treatment types also shifted through time (Fig. 1b). *C. penicillatus* had higher average abundance in the kangaroo rat exclosure plots before *C. baileyi* arrived. During the time *C. baileyi* was established, however, *C. penicillatus* had an even higher average abundance on controls. *C. penicillatus’s* preference for control plots increased with increases in *C. baileyi* abundance (Fig. 1c; y = −0.14x + 0.40, df = 16, RSE = 0.40, p < 0.001).

**Figure 1:**
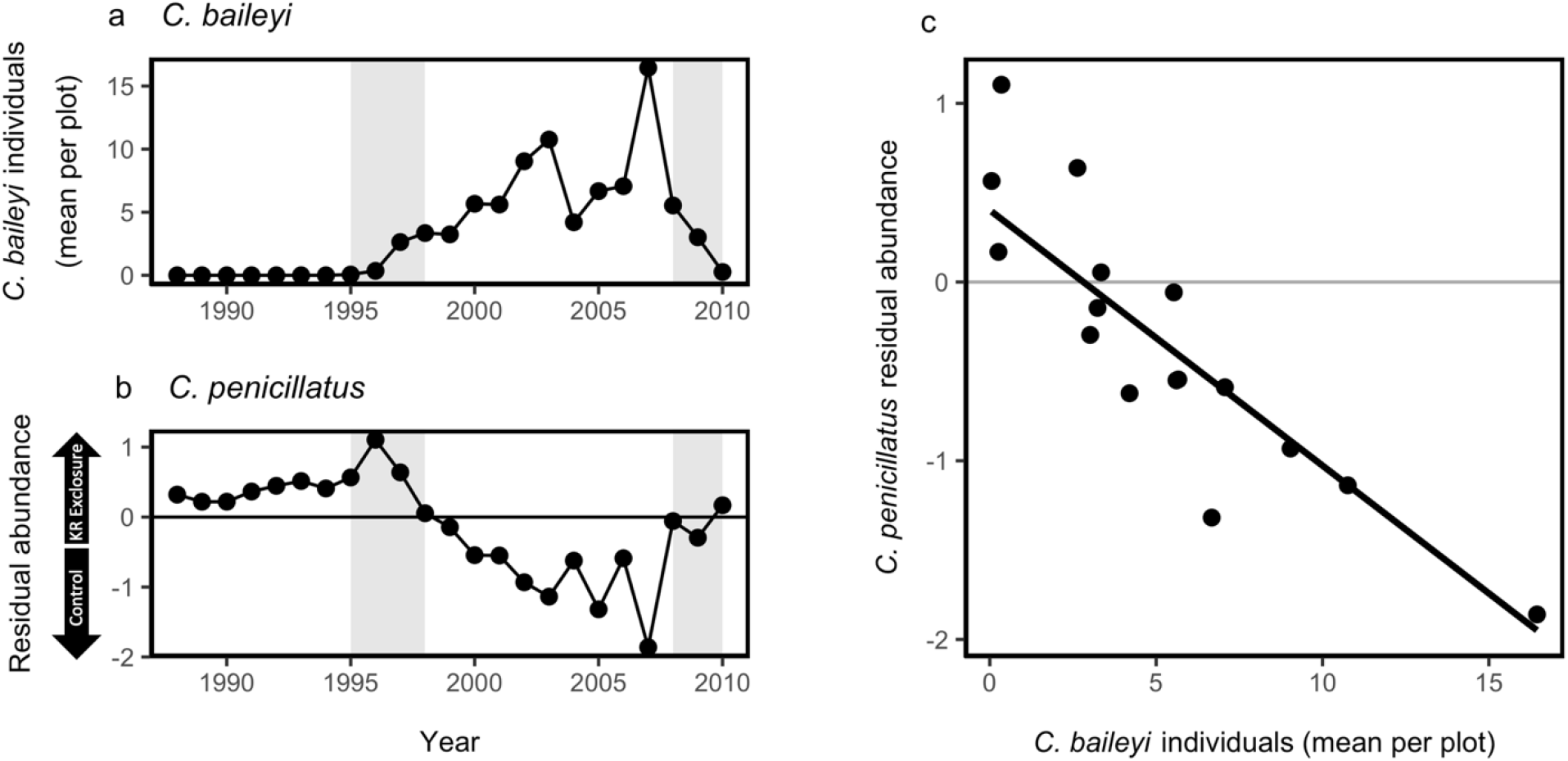
Relationship between *C. penicillatus* abundances on treatments and *C. baileyi* abundance. (a) Average number of *C. baileyi* individuals per plot through time. (b) The difference between mean *C. penicillatus* individuals per treatment through time. The zero line indicates equal numbers of *C. penicillatus* on both treatments. Points are residuals from a linear model run against a 1:1 line of mean *C. penicillatus* abundance on kangaroo rat exclosures against controls. Above the zero line (positive residuals) indicates higher mean *C. penicillatus* abundance on kangaroo rat exclosures; below the line (negative residuals) are higher mean *C. penicillatus* on controls. In (a) and (b), grey bars indicate the colonization period (1995-1998) and subsequent decline (2008-2010) of *C. baileyi*. (c) Generalized least squares regression of *C. penicillatus* differences from equal (y-axis from [a]) against mean *C. baileyi* individuals per plot per year (y-axis from [b]). As mean *C. baileyi* abundances increase, the mean abundance of *C. penicillatus* shifts from more individuals on kangaroo rat exclosures to more on controls.

### Population-level metrics of C. penicillatus

Residency of *C. penicillatus* depended on treatment and *C. baileyi* status (χ^2^ = 10.72, df = 3, p = 0.01). Before *C. baileyi* colonized the site, residency for *C. penicillatus* was significantly higher on kangaroo rat exclosures than on controls (Fig. 2a). This difference completely disappeared after *C. baileyi* established, at which point residency became statistically indistinguishable between treatments.

**Figure 2:**
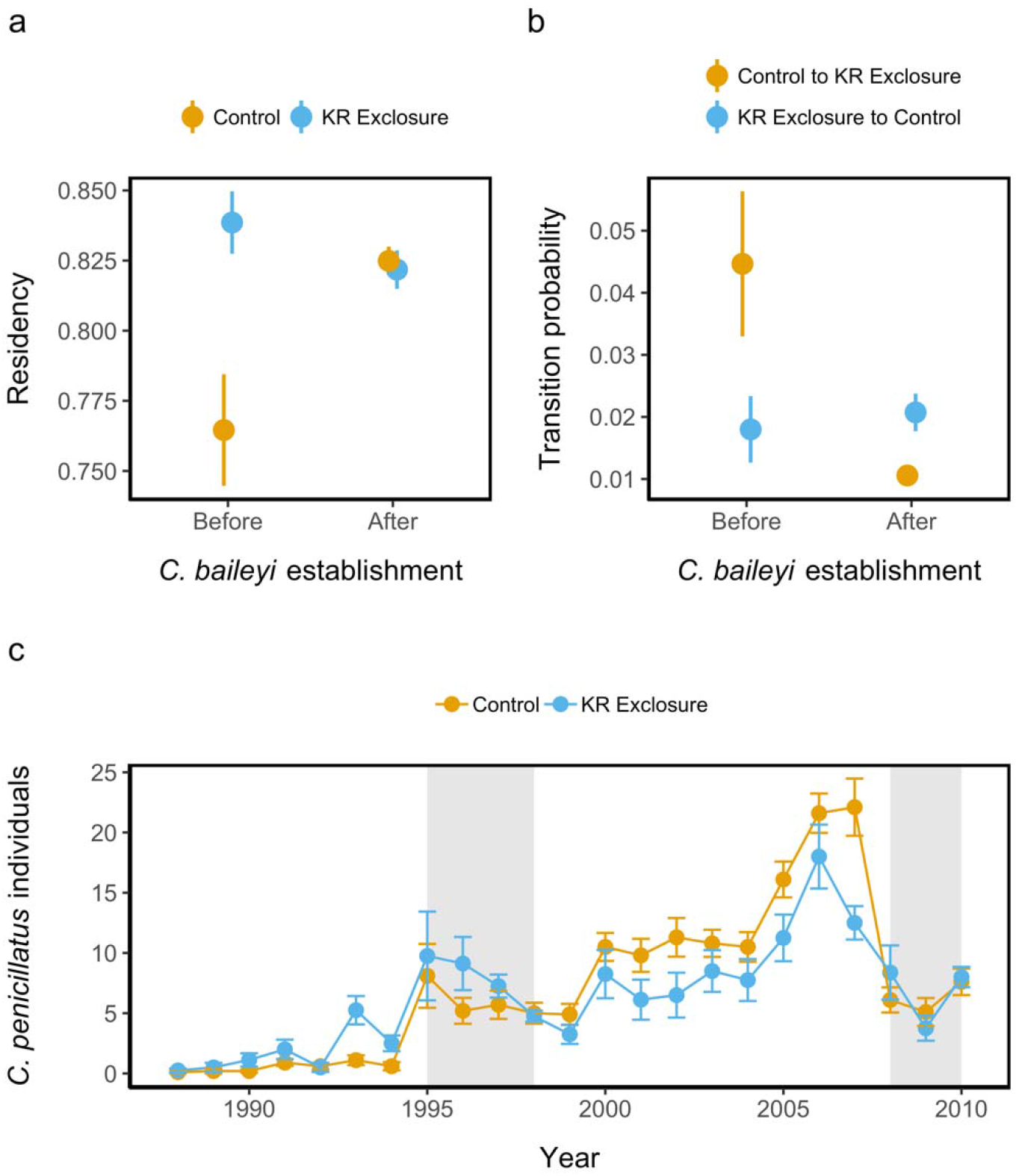
Population-level metrics of *C. penicillatus* by treatment type. (a) Residency of *C. penicillatus* by treatment and *C. baileyi* establishment in the system. (b) Probability of *C. penicillatus* individuals moving from one treatment to the other, also based on *C. baileyi* establishment. (c) Mean new *C. penicillatus* individuals per plot through time. Grey bars indicate the period of establishment (1995-1998) and subsequent decline (2008-2010) of *C. baileyi*.

The transition probability of *C. penicillatus* also depended on treatment and *C. baileyi* status (χ^2^ = 16.53, df = 3, p < 0.001). The probability of a *C. penicillatus* individual moving from a kangaroo rat exclosure to a control plot was low, regardless of *C. baileyi* establishment (Fig. 2b). When a *C. penicillatus* individual moved before *C. baileyi*’s arrival, it was more likely to move from a control plot to a kangaroo rat exclosure. Afterwards, however, the probability of a *C. penicillatus* individual moving from a control plot to a kangaroo rat exclosure was not only significantly lower than before *C. baileyi* establishment but also significantly lower than the probability of movement in the other direction (Fig. 2b).

The significant interaction between treatments and *C. baileyi* establishment in new (i.e., untagged) *C. penicillatus* individuals also supports changes in patch preference (Fig. 2c). Before the arrival of *C. baileyi*, kangaroo rat exclosures had significantly higher numbers of new individuals appearing (F_1,389_ = 24.87, p < 0.001); after *C. baileyi* established in the system, new individuals were consistently found on control plots in higher average numbers until the period of *C. baileyi* decline in the late 2000s.

### System-level aspects of patch preference

Prior to *C. baileyi* fully establishing in the system, the energy on kangaroo rat exclosures was only a fraction of that on control plots (Fig. 3). With the arrival of *C. baileyi*, however, the average energy on kangaroo rat exclosure plots reached over 80% of the energy used on control plots, even with *C. penicillatus* individuals moving to the control plots.

**Figure 3.**
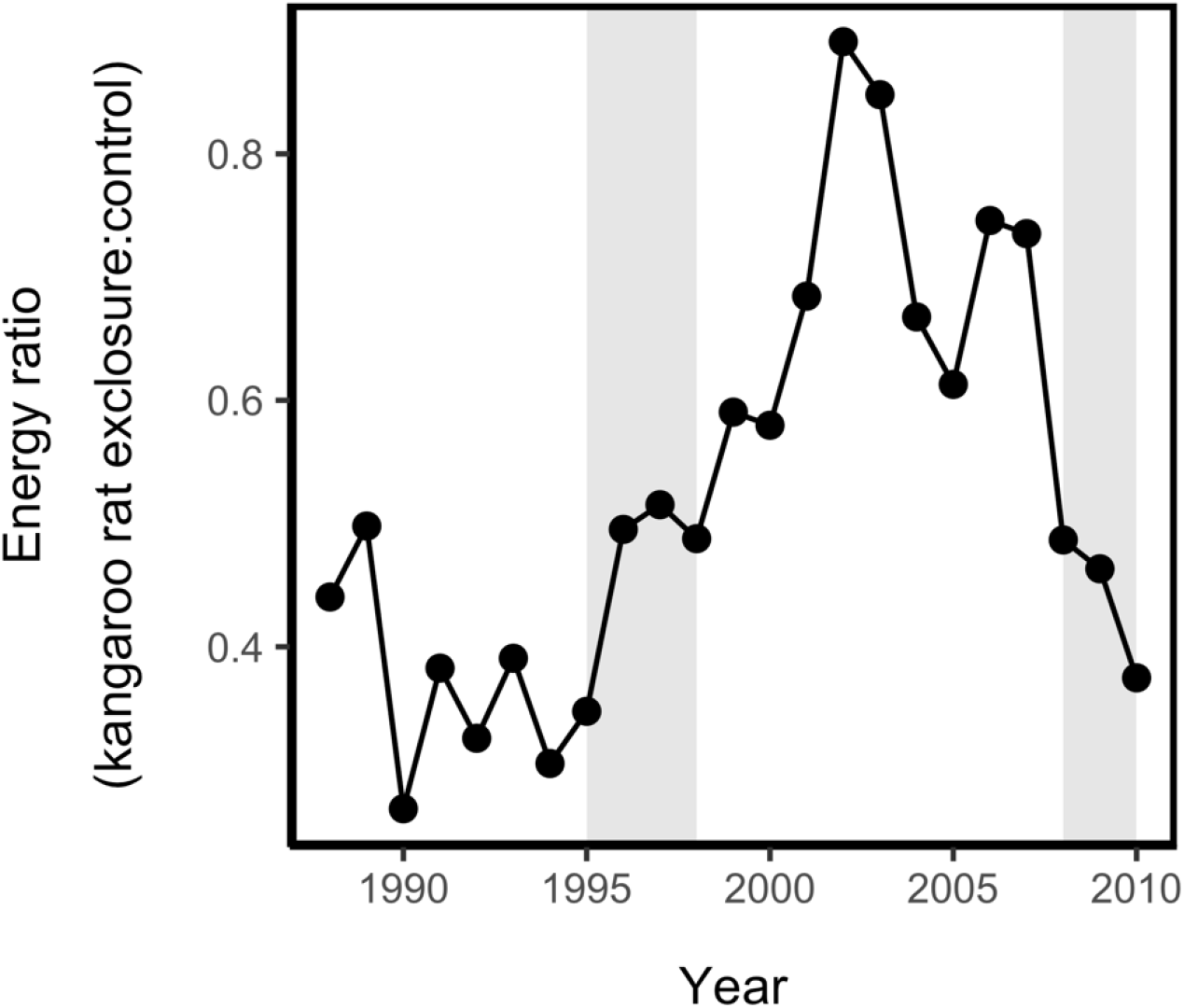
Ratio of total rodent energy in kangaroo rat exclosures to controls though time. Grey bars indicate establishment (1995-1998) and subsequent decline (2008-2010) of *C. baileyi*.

## Discussion

Our results suggest that the arrival of a congener changed perceptions of patch quality for one of the abundant rodent species in our ecosystem, *C. penicillatus*. Changes in patch use can be driven by perceived declines in an individual’s current patch or perceived increases in quality in another patch. Increases in new *C. penicillatus* individuals on the controls and declines in residency on kangaroo rat exclosures suggest that both of these mechanisms may have occurred with the arrival of *C. baileyi*, ultimately shifting patch use by *C. penicillatus*.

Declines in the perceived quality of kangaroo rat exclosures—as evidenced by declines in *C. penicillatus* residency—could have been caused by changes in resource availability after the arrival of *C. baileyi* (Fig. 1). Before *C. baileyi* established, rodent energy use on the kangaroo rat exclosures was never more than half of that found in the control plot (Fig. 3). With the preferential establishment of *C. baileyi* on the kangaroo rat exclosures, however, the energy ratio between the exclosure and control plots increased considerably. If patches have roughly the same amount of resources, patches with lower rates of energy use should have more resources that are not being fully exploited (MacArthur & Pianka 1966). Because the two species are congeners, niche similarities due to a shared evolutionary history may have increased the possibility of substantive overlap in their resource preferences. Thus, the colonization of *C. baileyi* on kangaroo rat exclosures may have had a substantive effect on rates of resource use on those plots, thus altering foraging and fitness expectations for *C. penicillatus*. This could explain the corresponding declines in residency for *C. penicillatus* (Fig. 2a) on kangaroo rat exclosures. However, while this explains why kangaroo rat exclosures may have been perceived as worse environments after the arrival of C. baileyi, this does not explain why controls were suddenly able to support higher numbers of *C. penicillatus*.

Greater increases in abundances of *C. penicillatus* on controls than on kangaroo rat exclosures suggest that *C. penicillatus* perceived improvements in the quality of those patches. Since *C. penicillatus’* residency and transition probabilities were similar between treatments after the arrival of *C. baileyi*, increases in abundance appear to be due to higher numbers of new individuals arriving on controls (Fig. 2c). These new individuals could be due to increases in birth rates, immigration rates or juvenile survivorship or decreases in juvenile dispersal rates, but we unfortunately do not have the data to discern the source of the new individuals.

Higher immigration rates into some patches over others can reflect differences in distances to source populations (i.e., mass effects or source-sink dynamics; Holt 1985, Pulliam 1988) or active decisions by individuals based on their expected fitness or resource intake rate in a patch, which is indicative of density-dependent habitat selection (e.g., Grant 1971, Brown & Munger 1985, Morris 1987, Abramsky et al. 1992, Morris & MacEachern 2010). Because our patches are interspersed in a matrix that is suitable habitat for *C. penicillatus*, all patches should be equidistant from a *C. penicillatus* source population. We also see no reason *C. penicillatus* should perceive control plots as improved based on abiotic conditions, habitat structure, or resource availability. At the scale of the site, all plots experience the same weather and our measure of rodent energy use suggests that resource availability on the two plot types is very similar (Fig 3). Furthermore, while the site has experienced vegetation changes (Brown 1998), there is no indication that this has differed by treatment (Ernest 2001).

The movement of *C. penicillatus* to control plots after the establishment of *C. baileyi* presents an interesting dilemma. If there were ecological opportunities (e.g., resource availability, territory, etc.) for *C. penicillatus* in the control plots after *C. bailey* become established on the kangaroo rat exclosures, why was *C. penicillatus* not utilizing that space previously? After *C. baileyi* established in the system, control plots tended to have a higher abundance of competitors (*Dipodomys spp*. and *C. baileyi)* than kangaroo rat exclosures (p < 0.001, Appendix S1: Fig. S5), making density-dependent habitat selection an unlikely explanation. Coexistence theory provides another interesting possibility. Resource competitors that would be unable to coexist if they interacted only with each other can actually benefit each other when they share a common competitor (Levine 1976, Stone & Roberts 1991, Allesina and Levine 2011). At our site, while *C. baileyi* showed a preference for kangaroo rat exclosures over controls, they were still present on control plots in considerable numbers (Fig. S1; Thibault and Brown 2008). Both natural history and observed dynamics at our site have shown that *C. baileyi* also competes with kangaroo rats (Thibault and Brown 2008), probably due to its similar size and dietary overlap (Reichman 1975). Thus, in this “enemy of my enemy is my friend” scenario, the shifts in the competitive network caused by adding *C. baileyi* to controls may have paradoxically improved conditions on control plots for *C. penicillatus*, leading to higher vital rates, more new individuals, and higher abundances on controls.

Species’ perceptions of patch quality can vary depending on a variety of factors, such as resource availability (MacArthur & Pianka 1966), biotic interactions (Grant 1971, Abramsky et al. 1992), and other habitat properties (Brown 1988). Changes in patch quality and selection can also affect communities and metacommunities through landscape-level processes (e.g., dispersal, colonization/extinction; Pulliam & Danielson 1991, Resetarits & Silberbush 2016). In this study, we used an experimental long-term study to show how species invasion and resulting shifts in the species composition can affect a species’ perception of patch quality and patch preference. This is not to suggest that changes in structural habitat or abiotic factors do not impact patch preference; much work in landscape ecology and metacommunity theory has shown that they can (Leibold & Chase 2018); rather, we use time series and experimentally manipulated patches to tease apart the effects of species composition from those of structural or abiotic habitat differences, changes which frequently occur together spatially. This method allows us to still assess spatial use patterns—a key aspect of metacommunity theory—while also allowing changes through time to inform our observations. We suggest that time is a key component in any holistic study of patch preference in community structure and metacommunity dynamics.

## Acknowledgments

We thank the UF Weecology group for helpful feedback, Dr. J. Simonis for statistical advice, and Dr. S. R. Supp for making R code from a previous project openly available, making our analyses substantially easier. We also thank Drs. D. Morris and B. Danielson and an anonymous reviewer for comments improving the manuscript. We thank the hundreds of people who have worked on the Portal Project to make these data available. This work was supported by National Science Foundation grant DEB-1622425 to SKME. EKB was also supported by the UF School of Natural Resources and Environment.

