## Appendix S1 for "Temporal changes in species composition affect a ubiquitous species’ use of habitat patches"

*E. K. Bledsoe and S. K. Morgan Ernest*

*Temporal changes in species composition affect a ubiquitous species’ views of patch quality*

*Ecology*

**Results with Data from 1989-2014:**

*Patch preference of C. penicillatus in response to C. baileyi abundance*

In addition to the linear generalized least squares model, we also fit a similar model with a first-order autoregressive structure included. In comparing the models with and without an autoregressive component, we found that they were not significantly different (χ^2^(1) = 1.66, *p* = 0.20). We chose to use the original model without an autoregressive structure as it was the most parsimonious model and the AIC values were nearly identical (no-correlation model: AIC = 36.86; correlation model, AIC: 37.20).

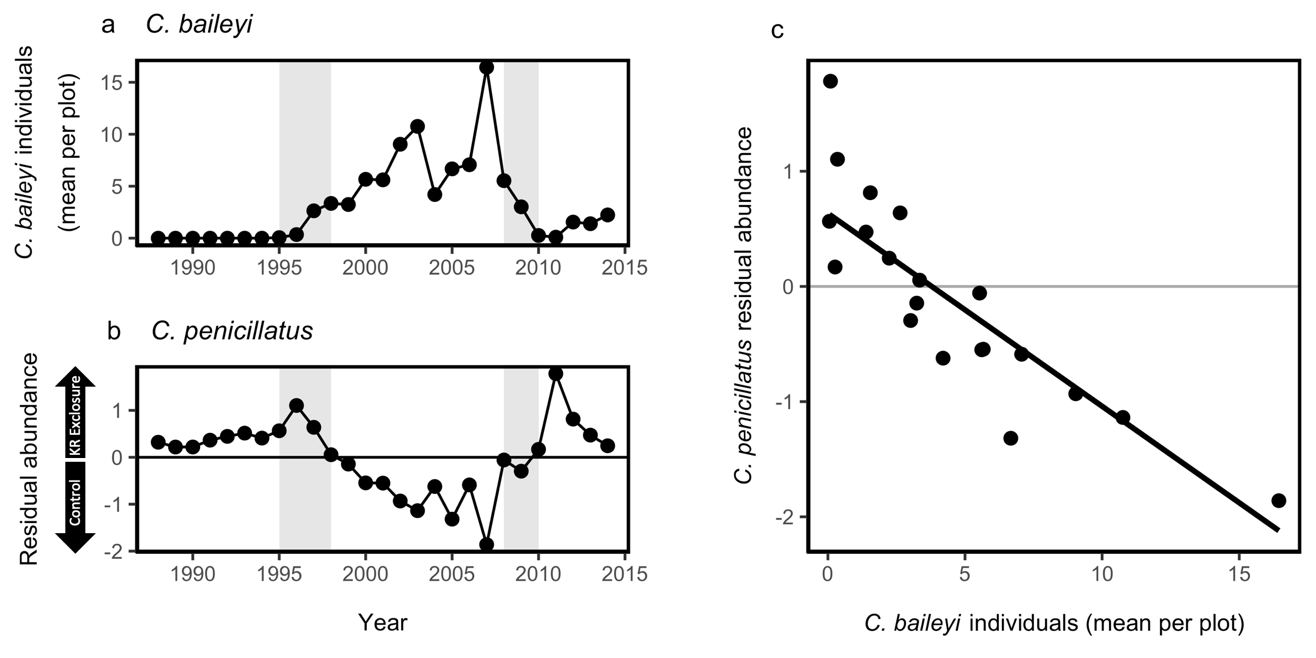

**Figure S1:** Relationship between *C. penicillatus* abundances on treatments and *C. baileyi* abundance. (a) Mean number of *C. baileyi* individuals per plot through time. (b) The difference between mean *C. penicillatus* individuals per treatment through time. The zero line indicates equal numbers of *C. penicillatus* on both treatments. Points are residuals from a linear model run against a 1:1 line of mean *C. penicillatus* abundance on kangaroo rat exclosures against control plots. Above the zero line (positive residuals) indicates higher mean *C. penicillatus* abundance on kangaroo rat exclosures; below the line (negative residuals) are higher mean *C. penicillatus* on control plots. In (a) and (b), grey bars indicate the colonization period (1995-1998) and subsequent decline (2008-2010) of *C. baileyi.* (c) Generalized least squares regression of *C. penicillatus* differences from equal (y-axis from (a)) against mean *C. baileyi* individuals per plot per year (y-axis from (b); y = -0.185x + 0.714, df = 20, RSE = 0.45, p <0.001). As mean *C. baileyi* abundances increase, the mean abundance of *C. penicillatus* shifts from more individuals on kangaroo rat exclosures to more on control plots.

Residency of *C. penicillatus* showed significant differences between treatment types and *C. baileyi* status (χ^2^ = 15.22, df = 3, p < 0.05). The transition probability of *C. penicillatus* also showed significant differences between treatment types and *C. baileyi* status (χ^2^ = 12.44, df = 3, p < 0.05).

| 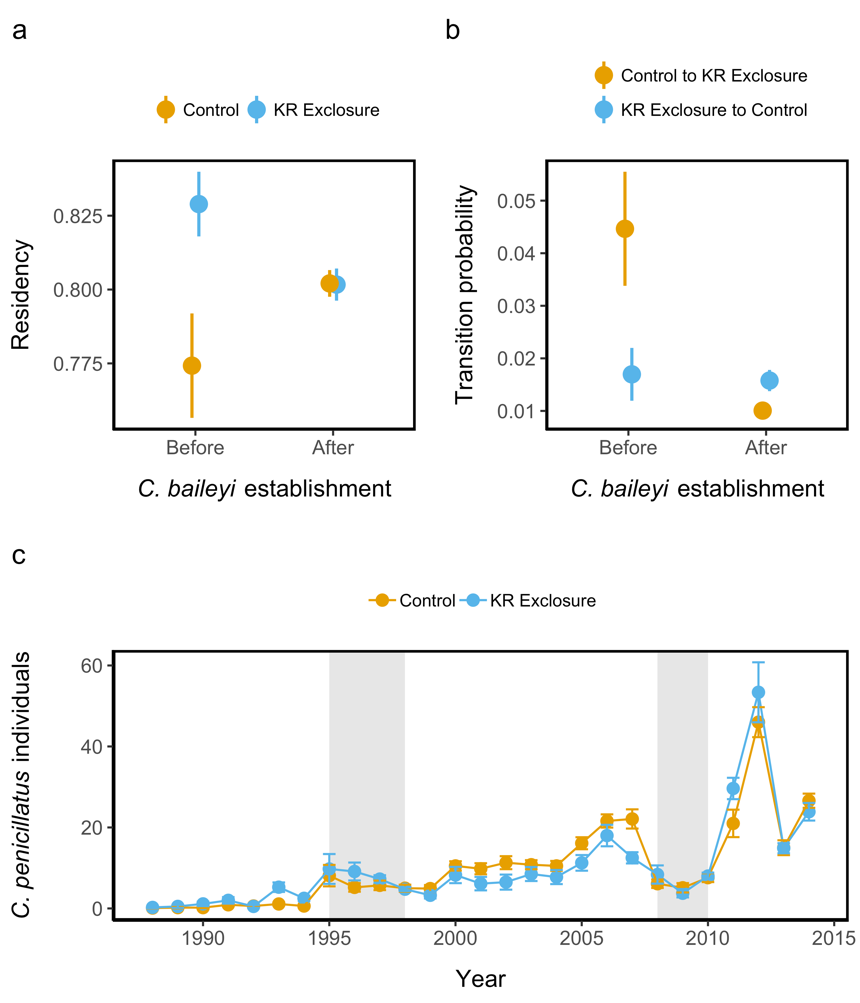 | **Figure S2.** Population-level metrics of *C. penicillatus* by treatment type. (a) Residency of *C. penicillatus* by treatment type and *C. baileyi* establishment in the system. (b) Probability of *C. penicillatus* individuals moving from one treatment type to the other, also based on *C. baileyi* establishment. (c) Mean new *C. penicillatus* individuals per plot through time. Grey bars indicate the period of establishment (1995-1998) and subsequent decline (2008-2010) of *C. baileyi.* Significantly more new individuals were caught on control plots after *C. baileyi* establishment in the system than on kangaroo rat exclosures (interaction term for treatment and time: p < 0.01). |
| --- | --- |

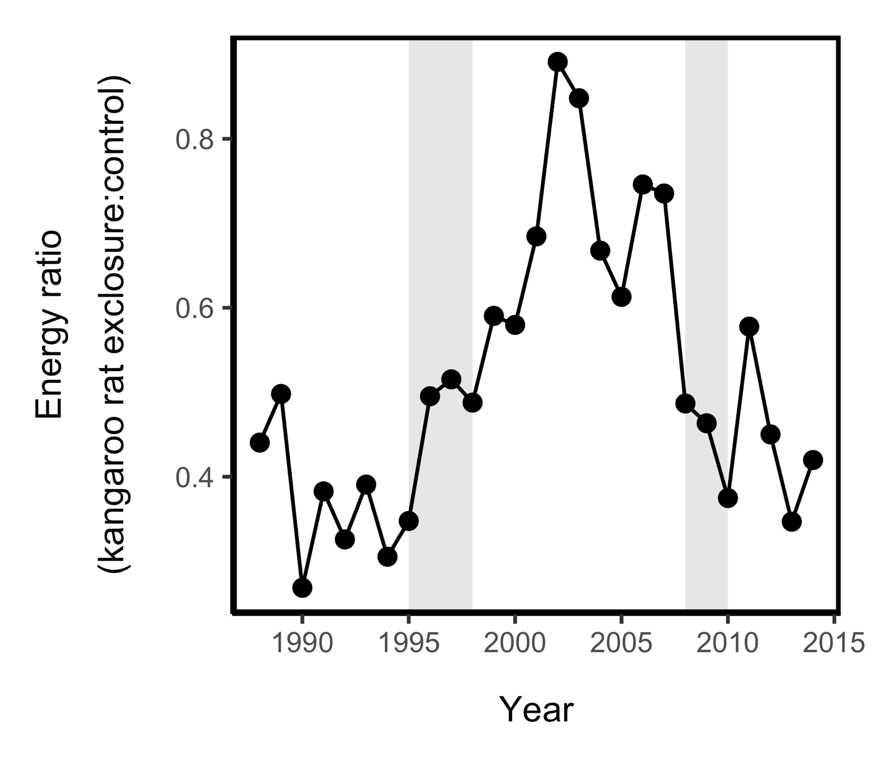

**Figure S3.** Ratio of total rodent energy in kangaroo rat exclosures to control plots though time. Grey bars indicate establishment (1995-1998) and subsequent decline (2008-2010) of *C. baileyi.*

**Additional Methods (1989-2010):**

***Study System and Data***

To calculate population metrics that required mark-recapture information on unique individuals, we performed extensive quality control and cleaning of the data to address potential issues (e.g., duplicate tags, uncertain species identification). Where individuals with identical tags could be determined to be unique individuals (based on time between capture or different species identifications), each individual was assigned a unique tag number for analysis. Those considered indeterminate were excluded from analysis.

For most analyses, we only used data from the controls and kangaroo rat exclosures. However, data from the rodent removal plots were incorporated in the RMark models to ensure the model estimates (estimated survival, transition probability) were as accurate as possible.

*The Curious Case of Chaetodipus baileyi*

We define *C. baileyi’s* arrival in the system as the first trapping event in which a *C. baileyi* individual was recorded at the site (September 1995), and we define *C. baileyi* establishment at the site as the first trapping event in which at least one *C. baileyi* individual was trapped on all eight kangaroo rat exclosure plots (July 1997). The period of time shown as the initial *C. baileyi* decline in the figures is the last trapping event in which *C. baileyi* was caught on all eight kangaroo rat exclosure plots (October 2008) and the first trapping event in which no *C. baileyi* individuals were caught throughout the site (November 2010).

The first *C. baileyi* was captured at the site in 1995, but the species remained relatively rare until 1999, when a large sheet flood at the site resulted in high mortality for kangaroo rat species but little to no mortality for *Chaetodipus* species. Although kangaroo rat abundances returned to previous levels within a few months, relative abundances of both *C. baileyi* and *C. penicillatus* remained higher than expected for nearly a decade (Fig. S4; Thibault & Brown 2008). The reduction in the competitively-dominant kangaroo rats—albeit temporarily—likely opened a space for *C. baileyi* to establish in the system and fill more ecological space where competitively-dominant kangaroo rats had once been (Brown & Heske 1990, Thibault & Brown 2008).

In 2009-2010, our site experienced strong drought conditions, and *C. baileyi* is known to be a drought intolerant species (Wilson & Ruff 1999). Although *C. baileyi* was already experiencing a population decline in 2008, the severe drought conditions almost completely extirpated the species from our site. On the other hand, the drought did not affect the *Dipodomys* (kangaroo rat) species as strongly. Without the reduction in competition from the kangaroo rat species, *C. baileyi* has not been able to re-establish at its previous abundance. While *C. baileyi* individuals continue to be trapped, the species is no longer a dominant species on any plot type.

***Analyses***

*Patch preference of C. penicillatus in response to C. baileyi abundance*

We fit linear generalized least squares models both with and without autoregressive structure included. In comparing the models, we found that the model with a first-order autoregressive structure fit the data significantly better than the model without (χ^2^_(1)_ = 4.79, *p* = 0.03).

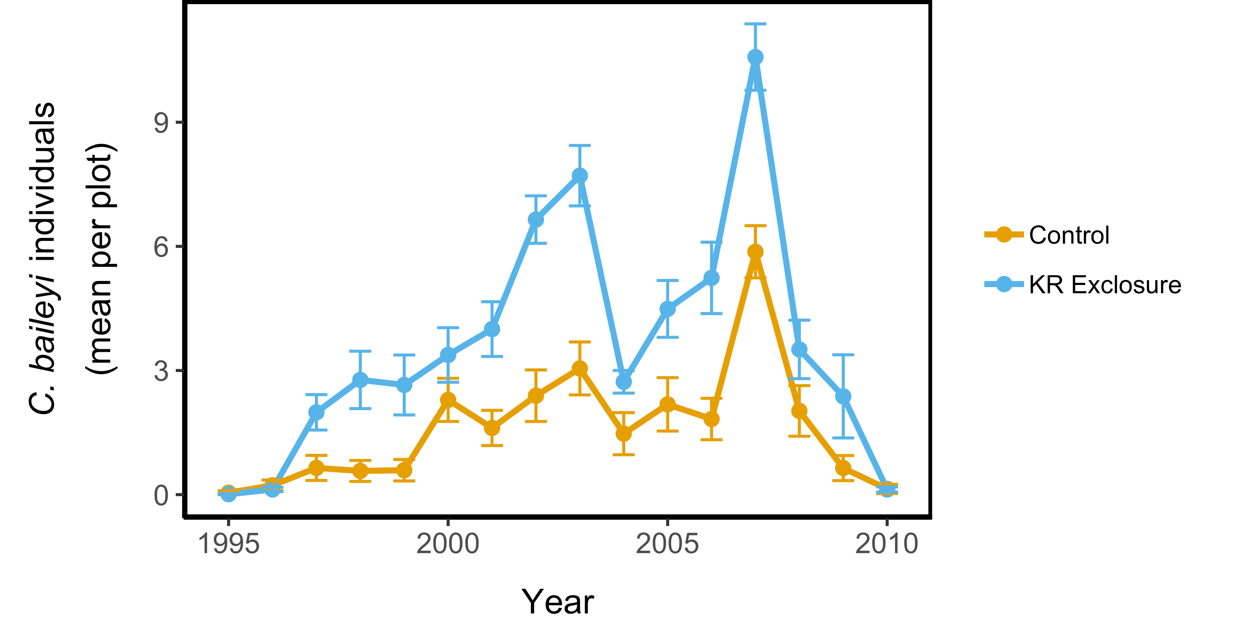

**Figure S4**. Mean (±SE) *C. baileyi* individuals per plot through time by treatment type. In most years, *C. baileyi* was found at higher frequencies in the kangaroo rat exclosures than on the control plots.

**Discussion:**

For density-dependent habitat selection to explain *C. penicillatus’s* change in habitat use to controls after *C. baileyi* entered the system, the number of competitors competing with *C. penicillatus* (*Dipodomys spp.* and *C. baileyi* individuals) would need to be higher on kangaroo rat exclosures. This is not the case, however. To test for density-dependent habitat selection after we used a linear mixed-effects model with year as a random effect to assess whether density-dependent habitat selection could explain the shift in *C. penicillatus’* habitat use after *C. baileyi* had established in the system (1998-2010). Post-*C. baileyi* establishment, the mean number of competitors differed significantly by treatment (F_(1, 220)_ = 16.80, p < 0.001), with control plots often having more competitors than kangaroo rat exclosures, indicating that density-depended habitat selection is not responsible for the shift in habitat use.

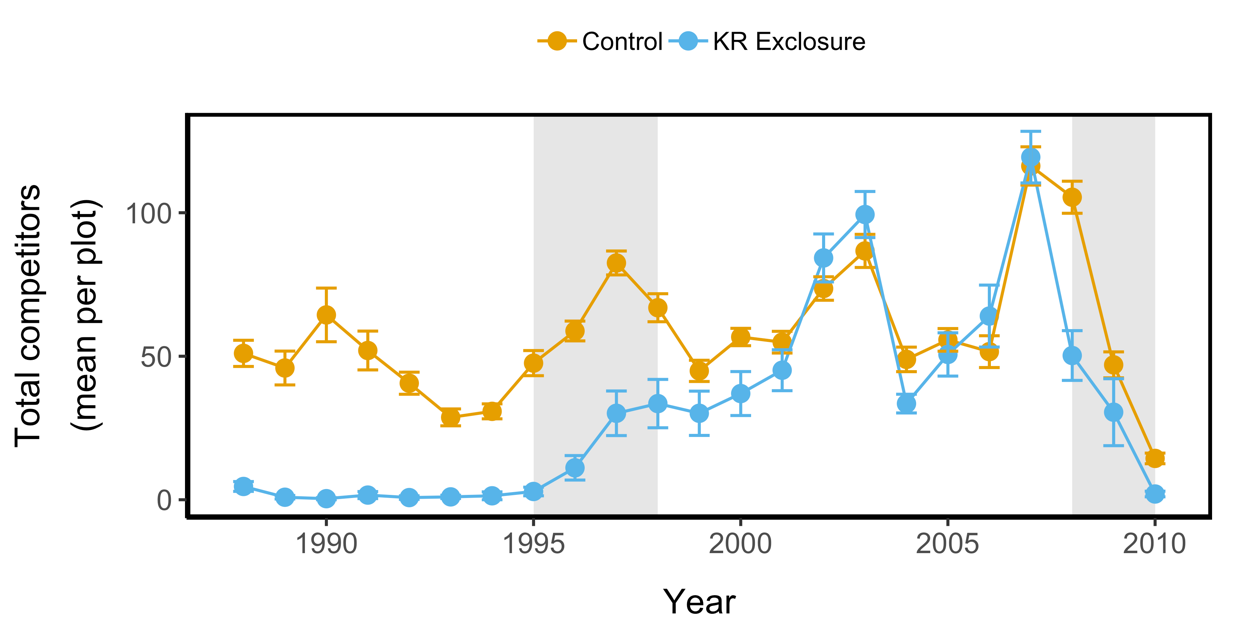

**Figure S5**. Mean (±SE) Mean number of *C. penicillatus* competitors (*Dipodomys spp.* and *C. baileyi*) individuals per plot through by treatment. After *C. baileyi* enters the system, the mean number of competitors differs between treatments (p < 0.001), with controls often having more competitors.
